# A fully phased accurate assembly of an individual human genome

**DOI:** 10.1101/855049

**Authors:** David Porubsky, Peter Ebert, Peter A. Audano, Mitchell R. Vollger, William T. Harvey, Katherine M. Munson, Melanie Sorensen, Arvis Sulovari, Marina Haukness, Maryam Ghareghani, Human Genome Structural Variation Consortium, Peter M. Lansdorp, Benedict Paten, Scott E. Devine, Ashley D. Sanders, Charles Lee, Mark J.P. Chaisson, Jan O. Korbel, Evan E. Eichler, Tobias Marschall

## Abstract

The prevailing genome assembly paradigm is to produce consensus sequences that “collapse” parental haplotypes into a consensus sequence. Here, we leverage the chromosome-wide phasing and scaffolding capabilities of single-cell strand sequencing (Strand-seq)^1,2^ and combine them with high-fidelity (HiFi) long sequencing reads^3^, in a novel reference-free workflow for diploid *de novo* genome assembly. Employing this strategy, we produce completely phased *de novo* genome assemblies separately for each haplotype of a single individual of Puerto Rican origin (HG00733) in the absence of parental data. The assemblies are accurate (QV > 40), highly contiguous (contig N50 > 25 Mbp) with low switch error rates (0.4%) providing fully phased single-nucleotide variants (SNVs), indels, and structural variants (SVs). A comparison of Oxford Nanopore and PacBio phased assemblies identifies 150 regions that are preferential sites of contig breaks irrespective of sequencing technology or phasing algorithms.

---

The first attempt to assemble a diploid human genome from a single individual (Craig Venter) capitalized on highly accurate and moderately long (500-1000 bp) Sanger sequencing reads^4^. However, such assemblies were highly fragmented and unable to resolve many repetitive regions of the human genome^4^. With the recent advances in long-read sequencing technologies (Pacific Biosciences of California, Inc. [PacBio] and Oxford Nanopore Technologies [ONT]), we are able to generate accurate and much more contiguous genome assemblies. By circumventing the problem of haplotype separation by sequencing fully homozygous hydatidiform mole cell lines^5, 6^ one can achieve highly contiguous assemblies, which in some instances traverse centromeric regions^7^. For diploid samples, local haplotype separation was previously demonstrated using long reads^8^ or linked reads^9^, but such approaches lack global phase information and are hence unable to separate haplotypes over extended genomic distances. Global haplotype partitioning of reads prior to assembly was shown to be possible by using trio data in conjunction with long reads. An approach that leverages parent-specific k-mers for this has been pioneered by Koren et al.^10^ However, such parental sequencing data are not always available, especially in clinical settings. Combining Hi-C data with long reads towards single-individual phased assembly has shown considerable promise^11,12^, but reliable scaffolding and phasing across the entire chromosomes still remains challenging.

Strand-seq is a single-cell sequencing method able to preserve structural contiguity of individual homologs in every single cell (**Fig. 1a**). This is achieved by utilizing a thymidine analog to selectively label and remove one of the DNA strands (the nascent strand, synthesized during DNA replication), which generates directional sequencing libraries of DNA template strands only (**Supplementary Notes**)^1,2^. Strand-seq comes with three important abilities: i) it can sort reads or contigs by chromosome^13–16^; ii) it can order and orient contigs^13^, and iii) it provides a chromosome-wide phase signal irrespective of physical distance^17^. Taken together, these features make Strand-seq the method of choice to be combined with high-accuracy long-read sequencing platforms to physically phase and assemble diploid genomes. This technique is particularly powerful when combined with other data types like linked reads or long reads to create dense long-range haplotypes^18^. Previously, we used this approach for partitioning reads prior to local assembly to improve structural variation sensitivity^19^ but read partitioning required mapping to a reference genome as an intermediate step, which can entail biases towards reference alleles and alignment artifacts. Here, we show how this limitation can be removed by exploiting Strand-seq’s additional ability to assign contigs to chromosomes in order to phase them and how this linking technology can be coupled with recent advances in highly accurate long-read sequencing. We present a completely reference-free workflow for diploid genome assembly where both parental haplotypes are accurately assembled into a ~6 Gbp genome.

**Figure 1:**
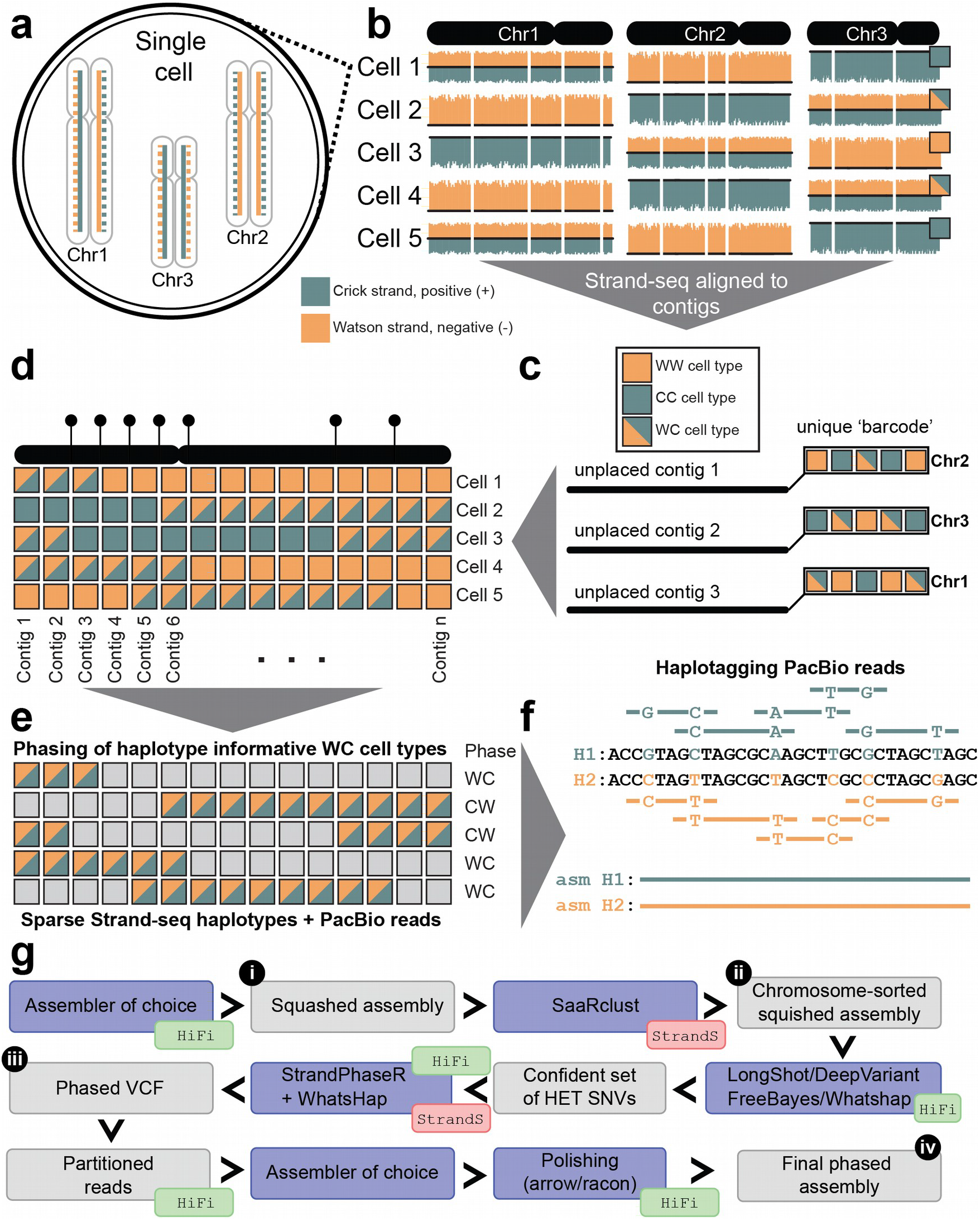
Conceptual overview of the complete genome assembly pipeline. **a**) In a single Strand-seq library only the template DNA strand (solid line) is sequenced for each parental homologous chromosome. **b**) Template strands of each homologue in a given diploid cell are randomly inherited by daughter cells (‘+’ positive strand, teal - Crick and ‘-’ negative strand, orange - Watson), resulting in three possible template-strand states for homologous chromosomes (height of bars plotted along each chromosome represents the number of ‘+’ and ‘-’ reads mapped in each genomic bin): WC - one Crick and one Watson strand represented for given homologues; WW - only Watson template strands represented; or CC - only Crick template strands represented. **c**) Unassigned contigs follow the same pattern of template-strand-state inheritance based on the homologue they belong to. **d**) Contig order can be inferred based on low frequency changes in a template-strand state resulting from sister chromatid exchange events (SCEs) in the parental cell: Contigs that are closer to each other tend to share the same template-strand state more often than more distant contigs. **e**) Regions with WC strand-state are haplotype informative and can be assembled into continuous haplotypes. **f**) Haplotypes can then be used to split long reads into their respective homologues. **g**) Generation of long-read (HiFi/CLR) based assemblies: i) Producing collapsed assemblies; ii) Assigning contigs to clusters using Strand-seq (StrandS); iii) Phasing clustered assemblies using the combination of Strand-seq and long PacBio reads; iv) Partitioning and reassembling of haplotype-specific PacBio reads and polishing of the final diploid assemblies.

Our unified assembly workflow starts by producing “collapsed” *de novo* assemblies from the full set of long reads from both haplotypes. We then align Strand-seq data to the contigs resulting from the *de novo* assembly (**Fig. 1b**). We use the SaaRclust package^15^, extended here to work with raw contigs (**Supplementary Notes**), to assign each contig to its respective chromosome (cluster) (**Fig. 1c**) and to infer the order of contigs within each chromosomal cluster by leveraging sister chromatid exchange (SCE) events identified with Strand-seq (**Fig. 1d**)^1,20,21^. This clustering by chromosome is a key step that enables haplotype phasing. To this end, we align both long single-molecule sequencing reads and Strand-seq data back to the clustered assemblies. Our assembly pipeline next calls heterozygous single-nucleotide variants (SNVs) using the long reads in order to obtain a confident set of markers for phasing. We use these heterozygous SNVs to reconstruct global chromosome-length haplotypes using WhatsHap^22,23^, combining Strand-seq and PacBio reads (**Fig. 1e**)^18^. The resulting phased SNVs are then used to tag and split long reads per haplotype, again using WhatsHap (**Fig. 1f**). For each set of haplotype-specific reads, our workflow performs a complete *de novo* assembly of each parental homolog, alternatively using wtdbg2^24^, Flye^25^, Canu^26^ or Peregrine^27^, and polishes the assemblies twice with Racon^28^ to obtain the final diploid assemblies (**Fig. 1g**).

To demonstrate the utility of our workflow for building a completely phased genome assembly, we generated ~33.4-fold HiFi sequence coverage from a single individual (HG00733) of Puerto Rican descent from the 1000 Genomes Project^29^ as well as ~32-fold and ~21-fold coverage of HiFi reads of the parental genomes (HG00731, HG00732) for validation purposes, respectively. We initially assembled HiFi reads for HG00733 using Canu^26^, into a haplotype-unaware (“collapsed”) assembly with contig N50 values of 14.9 Mbp. To scaffold the genome, we aligned 115 single-cell Strand-seq libraries generated for HG00733 in the context of the Human Genome Structural Variation Consortium (HGSVC)^19^ to the collapsed assembly. The cumulative depth of Strand-seq reads was 2.87-fold and covered 73% of genomic positions in the assembly. After clustering collapsed contigs by chromosomes using SaaRclust, we aligned all contigs back to GRCh38 for evaluation purposes. Overall, ~78% mapped back to their respective chromosome of origin, with the bulk of misassignments corresponding to small contigs (mean size 174,807 bp). Importantly, ~99.6% of the total length of all clustered contigs were assigned to their correct chromosomal origin (**Fig. 2a**).

**Figure 2:**
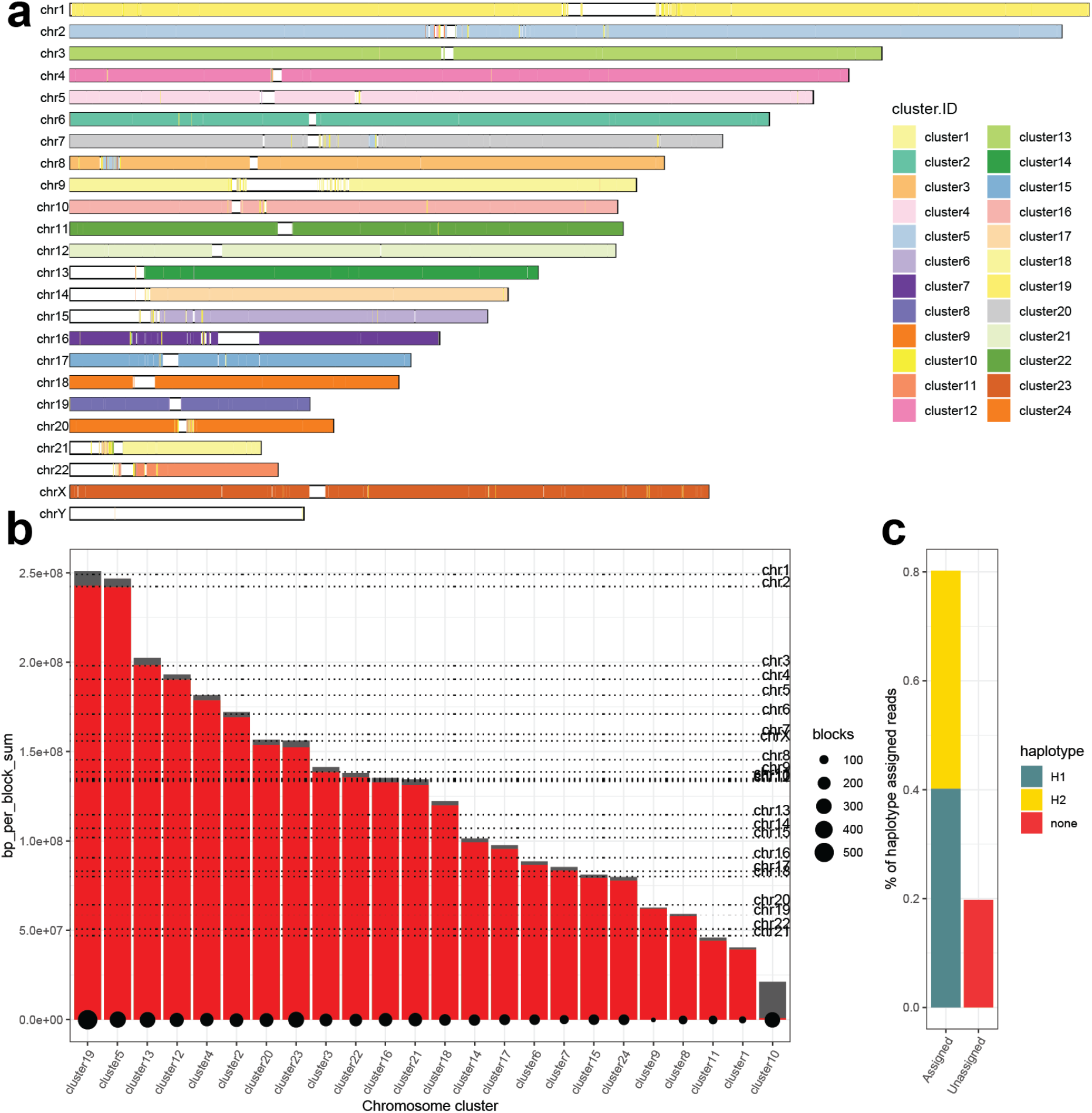
Reference-free scaffolding and phasing of collapsed assembly for HG00733. **a**) Each contig represents a range based on mapping coordinates on GRCh38. Contigs are colored based on cluster identity determined by SaaRclust. In an ideal scenario there is a single color for each chromosome. **b**) A barplot that shows the total length of all haplotype blocks per cluster in dark gray. The size of the longest haplotype block is shown on top in red. Size of the point at the bottom of each bar reflects the number of haplotype blocks in each cluster. For perspective, the real size of each chromosome for GRCh38 is plotted as a horizontal dotted line. **c**) The percentage of PacBio reads successfully assigned to either haplotype 1 (H1 - teal) or haplotype 2 (H2 - yellow). Reads that could not be assigned to either haplotype are shown in red.

Using DeepVariant, we detected 2,525,898 heterozygous SNVs genome-wide within the collapsed assembly. Phasing these variants using the Strand-seq signal and the HiFi reads^18^ resulted in chromosome-length haplotypes with >95% (**Supplementary Fig. 1,** red line) of all these heterozygous variants placed into a single haplotype block. Importantly, the longest haplotype block spanned almost the entire length of each cluster (red bars **Fig. 2b**) and closely matched the expected chromosome lengths from GRCh38 (dotted horizontal lines **Fig. 2b**). With such global and complete haplotypes we assigned ~81% of the original HiFi PacBio reads to either parental haplotype 1 (H1) or haplotype 2 (H2) (**Fig. 2c**). The remaining ~19% of haplotype-unassigned reads likely originate from autozygous regions and low mappability regions such as SDs and heterochromatic regions. As expected, we also found unassigned reads to be slightly shorter than haplotagged reads (**Supplementary Fig. 2**). These results are comparable with the previous study^3^ that used family trio information to haplotag 79.2% of HiFi PacBio reads.

We next assembled haplotype-specific reads into completely phased *de novo* assemblies using one of the most popular assemblers, Canu^26^, and the recently described fast assembler for HiFi data, Peregrine^27^. While Peregrine generated more contiguous genome assemblies (N50 contig: H1: 28 Mbp, H2: 29.1 Mbp) compared to Canu (H1: 9.9 Mbp, H2: 10.7 Mbp), we noted more misassemblies, especially chimeric contigs, near the end of contigs (**Supplementary Table 1**). We detected a total of 14 and 21 assembly errors in Canu and Peregrine phased assemblies, respectively (**Supplementary Table 1, Supplementary Notes**). Using Strand-seq data and SaaRclust, we readily corrected contig misorientations and chimerisms in the Peregrine assembly (**Supplementary Fig. 3**) and found the majority (~76%) of these misassemblies overlapped or mapped in the vicinity of SDs of size 50 kbp and longer (**Supplementary Fig. 4a,b**). This is expected as highly identical SDs promote false joins during the assembly process^30^. After correcting misassemblies, the final contig N50 remained high (H1: 25.8 Mbp and H2: 28.9 Mbp).

Using Strand-seq’s capacity to preserve structural and directional contiguity of individual homologs, we assigned phased Peregrine contigs into whole chromosomal scaffolds, again using the process described above for the collapsed assemblies. First, we assigned each contig to its chromosome of origin (**Supplementary Fig. 5a**), with more than 99.8% of a total contig length correctly assigned for both haplotype assemblies. Second, we synchronized the orientation of all contigs within each chromosomal scaffold in both haplotypes. Remarkably, after the contig reorientation process, 99.7% and 99.6% of a total contig length mapped to GRCh38 in a single direction for H1 and H2, respectively (**Supplementary Fig. 5b**). Lastly, we ordered contigs within both phased assemblies, obtaining an ordering that highly correlated (H1: 0.959, H2: 0.964) with the expected contig order as defined by mapping contigs to GRCh38 (**Supplementary Fig. 5c,d,e**).

To confirm the haplotype-resolved genome assemblies were correctly phased across all chromosomes, we independently assigned each 1 Mbp window of the assembled contigs to one of the two parents (*i.e*. HG00731, and HG00732; **Methods**) by using a set of trio-phased SNVs produced earlier^19^. As expected, the proband (HG00733) assembly was correctly phased, with only sporadic local errors (**Fig. 3a**) amounting to a switch error rate of 0.4%. To specifically assess phasing performance at a challenging but biomedically relevant locus, we examined the whole major histocompatibility complex (MHC) region and found that it was traversed by a single contig in both haplotype assemblies. These assemblies were phase consistent with recently released Shasta assemblies^31^ that used trio-binned ONT data, with a Hamming error rate of 0.28% (**Methods, Supplementary Fig. 6**) and represented some of the most diverse regions of the genomes (**Fig. 3b**).

**Figure 3:**
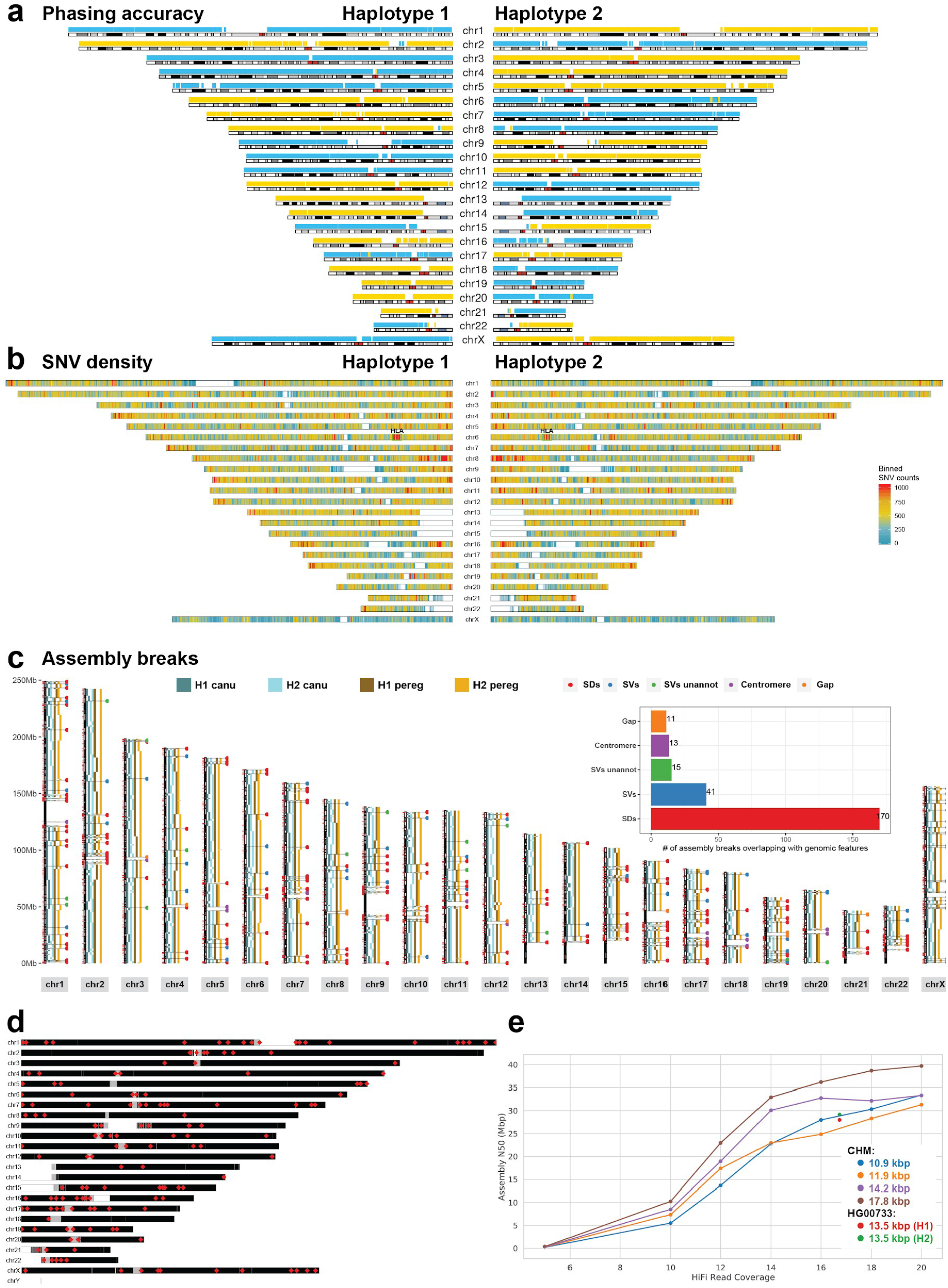
Phased assembly analysis and common assembly breaks. **a**) Each 1 Mbp block of phased contigs (Freeze 1.1, see Data Availability) are assigned to one of the parental genomes using SNV data from the parents^19^: maternal segments (HG00732) are shown in blue and paternal segments (HG00731) are shown in yellow. **b**) Genome-wide summary of SNV density counted in 500 kbp genomic bins sliding by 10 kbp. The HLA locus on Chromosome 6 is labeled as “HLA”. **c**) An ideogram that shows aligned contigs separately for haplotype 1 (H1) and haplotype 2 (H2). Subsequent contigs are plotted as discontiguous rectangles along each chromosome. Positions of common breaks (n = 250) between Canu and Peregrine assemblies are highlighted by horizontal lines and their overlap with various genomic features such as SDs are marked by colored dot. **Inset:** A barplot summarizing total counts of each genomic feature across all 314 assembly breaks. **d**) An ideogram that shows genomic positions of 150 common assembly breaks shared by multiple assembles. Gray rectangles represent centromeric positions while white rectangles points to genome gaps. **e**) Plots the effect of coverage and read length on assembly contiguity. Points connected by lines represent the N50s of Peregrine assemblies for CHM libraries as a function of coverage (blue, CHM13, 10.9 kbp; orange, CHM1, 11.9 kbp; purple, CHM13, 14.2 kbp; brown, CHM13; 17.8 kbp). These assemblies show what contiguity is attainable with Peregrine given different read lengths and coverages in a genome with only one haplotype. Highlighted in red and green are the two Peregrine assemblies of the haplotypes of HG00733 (red, H1, 13.5 kbp; green, H2, 13.5 kbp).

We generated estimates of the consensus quality value (QV) of our assembly using several independent methods. We sequenced and assembled 77 random BACs from an HG00733 clone library (VMRC62) and compared these sequences to the phased assemblies to estimate the consensus QV of the assembly (**Methods**). We find that the median QV of the random BACs aligned to the assembly to be 40.78, which corresponds to less than one error every 10,000 bases. Additionally, we derived QV estimates based on three variant call sets generated by mapping Illumina short reads (HG00733) and HiFi sequencing of parents (HG00731 and HG00732) to the assemblies. By identifying homozygous calls within high-confidence regions (**Methods**), we computed QV estimates ranging from almost 40 (parent HiFi reads) up to an upper bound of 60 (Illumina reads) (**Supplementary Table 2A/2B, Methods**). Overall, our QV estimates are comparable to the QV achieved in the HiFi assembly of a haploid human genome, CHM13 (e.g., BAC QV 40.78 vs. 45.25, Vollger et al., 2019). Despite the lower coverage per phased haplotype, we were able to resolve a comparable level of SDs on both haplotypes. We estimate that 33.9% and 34.5% of SDs were resolved in the H1 and H2 assemblies of HG0733, respectively (**Methods**). This estimate is similar to Peregrine assemblies of CHM13 assembled with 16- and 18-fold coverage—both of which resolved an estimated 35.8% of SDs. The H1 and H2 assemblies both showed signs of increased read coverage over 28.9 Mbp and 29.1 Mbp of their respective assemblies (**Methods**). These regions likely represent collapsed repetitive sequence that is unresolved in the *de novo* assembly of each haplotype. Of these regions, 166 (H1) and 162 (H2) correspond to collapsed duplicated regions greater than 50 kbp in length and are similar to unresolved regions observed in other haploid human genome assemblies (see below).

To discover genetic variation, we aligned contigs from both haplotypes to GRCh38 and identified SNVs, indels and structural variants (SVs) based on a previously described approach^30^, which were then merged to create a set of heterozygous and homozygous calls (**Methods**). From this analysis, we identified a total of 4.1 million SNVs (2.8 million per haplotype) (**Fig. 3b**) and 1.01 million indels distributed among insertions and deletions (515,224 and 494,810, respectively) (**Supplementary Table 3, Supplementary Fig. 7a**). Regions of increased genetic diversity were observed near the telomeres and HLA as expected (**Fig. 3b, Supplementary Fig. 7b,c**). We also identified SVs including 15,139 insertions and 9,579 deletions (**Supplementary Table 4, Supplementary Fig. 7b**). Considering gene-disruptive indels and SVs, we observe 198 disrupted genes in our diploid genome compared to 135 per haploid genome^32^ (**Supplementary Table 5**). If we exclude repetitive regions, where variants are often difficult to compare because of alignment issues, and use Human Genome Structural Variation Consortium (HGSVC) HG00733 calls^19^ as a truth set, we estimate 92% sensitivity and 92% specificity (**Supplementary Fig. 8**). If we include repetitive regions, we estimate 67% sensitivity and 75% specificity mostly due to a difficulty in comparing variant calls in tandem repeat sequences (**Supplementary Fig. 9**).

There are regions of the genome that have been notoriously difficult to assemble even with long-read technologies^6,33^. In this study, we operationally defined such difficult regions of the human genome as positions where both the phased Canu and Peregrine assemblies consistently break. In total we have localized 250 common breaks in our phased *de novo* assemblies (**Fig. 3d**). The majority (68%) of these assembly breakpoints lie within SD-rich regions of the genome that are copy number variable. Of these breaks, 41 correspond to previously detected SVs not associated with segmental duplication^19^. Notably, we also observed 15 assembly breaks where contiguous read-depth profiles suggest the presence of SVs that were missed as part of the HGSVC effort on this specific individual (**Supplementary Fig. 10**). To define if these 250 common assembly breaks are shared among other phased assemblies, we examined a recently released Shasta ONT assembly of the same individual^31^. We found that 150 of those breaks disrupt the Shasta assembly as well (**Fig. 3e, Supplementary Table 6**). Strikingly, we found 129 of these regions overlap SVs detected by the HGSVC, of which 76 were genotyped as inversions. This is expected as inversions are often flanked by highly identical SDs and are perhaps the most difficult class of SVs to detect and genotype. Interestingly, even the most contiguous telomere-to-telomere assembly of haploid CHM13 genome, constructed from ultra-long ONT reads and PacBio data, shares 64 common assembly breaks found in aforementioned assemblies. We propose that these universal assembly breaks (UAB) represent regions of our genome where neither the sequencing technology nor assembly algorithms have yet sufficiently evolved to resolve the underlying sequence in an automated fashion. These UAB regions represent compositional features of the human genome and not the result of incomplete phasing of long-read data. For example, even when sequence reads are fully phased (as in the case of haploid genomes), increasing coverage and insert size only moderately improves contiguity (**Fig. 3e**) and the two human genomes we assembled here have reached that empirical upper bound based on comparisons to human haploid references^32^.

In summary, we have introduced a novel assembly workflow to combine Strand-seq and long PacBio reads in a completely reference-free manner to provide fully phased and highly contiguous *de novo* assemblies of diploid human genomes. Previously, this was only possible by resorting to parental genome sequencing. Our assembly strategies allow us to transition from “collapsed” human assemblies of ~3 Gbp to fully phased assemblies of ~6 Gbp where all genetic variants, including SVs, are fully phased at the haplotype level. Even though we showcase our method using HiFi reads, the principle is applicable to other sequencing technologies, including continuous long-read (CLR) reads and ONT sequencing data. Our pipeline is designed to accommodate a range of assembly tools, including Canu^26^, Peregrine^27^, wtdbg2^24^ and Flye^25^, and different variant callers, including FreeBayes^34^, LongShot^35^, DeepVariant^36^, and WhatsHap genotyping^37^. This method should open the door to producing high-quality phased human genomes needed for personalized SV discovery in healthy and diseased individuals. Fully phased, reference-free genomes are also the first step in constructing comprehensive human pangenome references that aim to reflect the full range of human genome variation^38^. Importantly, our work also highlights recalcitrant regions of genome assembly irrespective of the approach and such challenging regions will require further technological and algorithmical advances moving forward.

## ONLINE METHODS

### Cell lines

Cell lines for Puerto Rican individuals HG00731, HG00732, and HG00733 have been previously described^19^.

### HiFi PacBio sequencing

Isolated DNA was prepared for HiFi library prep as described^3^. Briefly, DNA was sheared to an average size of about 15 kbp using Covaris gTUBE, and quantity and size checked using Qubit (ThermoFisher) and FEMTO Pulse (Agilent) instruments. Fragments underwent library preparation using the Template Prep Kit v1 (PacBio), then fractionation on a SageELF (Sage Science) instrument. After evaluating size, fractions averaging 11, 13, or 15 kbp were sequenced on a Sequel II (PacBio) instrument using Sequel II chemistry v1 or v2EA. After sequencing, raw data was analyzed with SMRT Link 7.1 or 8.0 using the circular consensus sequencing (CCS) protocol with a cutoff minimum of three passes and estimated accuracy of 0.99. In total, 18 SMRT Cell 8Ms were run for the Puerto Rican trio (HG00731, HG00732, HG00733) for an average yield per sample of 91 Gbp of HiFi reads (**Supplementary Table 7**).

### Strand-seq data analysis

All Strand-seq data in a FASTQ format have been obtained from publicly available sources (**Data availability**). At every step that requires alignment of short-read Strand-seq data to the collapsed or clustered *de novo* assembly (**Fig.1**), we used BWA-MEM (version 0.7.15-r1140) with the default parameters. In the next step we filtered out all secondary and supplementary alignments using SAMtools (version 1.9). Subsequently, duplicate reads were marked using Sambamba (version 0.6.8). At every Strand-seq data analysis we filtered out reads with mapping quality less than 10 as well as all duplicate reads.

### Collapsed genome assembly

Initially, collapsed assemblies were constructed in order to produce a set of unphased contigs. We assembled HiFi reads using the Canu and Peregrine assemblers.

All Peregrine (0.1.5.3) assemblies were run using the following command:

~~~
pg_run.py asm {reads.fofn} 24 24 24 24 24 24 24 24 24 --with-consensus \
       --shimmer-r 3 --best_n_ovlp 8 --output {assembly.dir}
~~~

All Canu (version v1.7.1) assemblies were run using the following command:

~~~
canu -d {assembly.dir} -p {assembly.prefix} genomeSize=3.1g \
       correctedErrorRate=0.015 ovlMerThreshold=75 \
       batOptions=“-eg 0.01 -eM 0.01 -dg 6 -db 6 -dr 1 -ca 50 -cp 5” \
       -pacbio-corrected $(cat {reads.fofn})
~~~

### Clustering contigs into chromosomal scaffolds

We used the R package SaaRclust^15^ to cluster *de novo* collapsed assemblies into chromosomal scaffolds. SaaRclust takes as an input Strand-seq reads aligned to the collapsed *de novo* assembly in a BAM format. Given the parameter settings, we discarded contigs shorter than 100 kbp from further analysis. Remaining contigs were partitioned into bins of size 100 kbp with 50 kbp overlaps. The counts of aligned reads per bin, separated by directionality (plus/Crick or minus/Watson), are used as an input for SaaRclust that divides contigs into a user-defined number of clusters (set to n = 100). Contigs genotyped as WC in the majority of cells were discarded. We further removed contigs that could be assigned to multiple clusters with probability p < 0.25 (**Supplementary Fig. 11**). Subsequently, SaaRclust merges clusters that share the same strand inheritance across multiple Strand-seq libraries. Shared strand inheritance is used to construct a graph of connected components (clusters) and the most connected subgraphs are reported, resulting in approximately 24 clusters, i.e., one cluster should ideally be representative of one human chromosome. Next, we defined misoriented contigs within each cluster as those having opposing directionality in every Strand-seq library. We used hierarchical clustering to detect groups of minus and plus oriented contigs. To synchronize contig directionality we switch direction in one group of contigs from plus to minus or vice versa. Contigs synchronized by direction are then subjected to positional ordering within a cluster. We again use contig strand-state coinheritance as a proxy to infer physical distance for each contig pair in every Strand-seq library. The resultant coinheritance matrix serves as input for the ‘Traveling Salesman Algorithm’ implemented in R package TSP (version 1.1-7)^39^ and attempts to order contigs based on strand-state coinheritance. As the initial collapsed assembly may contain assembly errors SaaRclust is able to detect and correct such errors as bins of the same contig being assigned to different clusters (‘Chimeric contig’) or bins of the same contig that differ in directionality (‘Misoriented contig’). Lastly, we export clustered, reoriented, and ordered contigs into a single FASTA file with a single FASTA record per cluster. A complete list of parameters used to run SaaRclust in this study is reported below:

SaaRclust command:

~~~
scaffoldDenovoAssembly(bamfolder = <>, outputfolder = <>, store.data.obj =
TRUE, reuse.data.obj = TRUE, pairedEndReads = TRUE, bin.size = 100000,
step.size=50000, prob.th=0.25, bin.method = ‘dynamic’, min.contig.size =
100000, assembly.fasta = assembly.fasta, concat.fasta = TRUE, num.clusters =
100, remove.always.WC = TRUE, mask.regions = FALSE)
~~~

SaaRclust command: (refining phased assemblies)

~~~
scaffoldDenovoAssembly(bamfolder = <>, outputfolder = <>, store.data.obj =
TRUE, reuse.data.obj = TRUE, pairedEndReads = TRUE, bin.size = 100000,
step.size = 100000, prob.th=0.9, bin.method = ‘dynamic’, ord.method =
‘greedy’, min.contig.size = 100000, min.region.to.order = 500000,
assembly.fasta = assembly.fasta, concat.fasta = FALSE, num.clusters = 100,
remove.always.WC = TRUE, mask.regions = FALSE)
~~~

### Variant calling

Clustered assemblies in full chromosomal scaffolds are then used for realignment of long PacBio reads. In order to call variants in HiFi PacBio reads, we use DeepVariant^40^ v0.8.0, which uses a deep neural network and a pretrained “pacbio_standard” model. For these variants, HiFi PacBio reads were aligned using pbmm2 v1.1.0 (https://github.com/PacificBiosciences/pbmm2) with the following settings: `“align --loglevel DEBUG --preset CCS --min-length 5000”` and filtered with `“samtools view -F 2308”`. After variant calling we select only heterozygous SNVs using BCFtools v1.9.

### Phasing chromosomal scaffolds

To create completely phased chromosomal scaffolds, we used a combination of Strand-seq and long-read phasing^18^. First, we realigned Strand-seq data on top of the clustered assemblies as stated previously. Only regions that inherit a Watson and Crick template strand from each parent are informative for phasing and are detected using breakpointR^41^ Haplotype-informative regions are then exported using breakpointR function called ‘exportRegions’. Using the set of haplotype-informative regions together with positions of heterozygous SNVs, we ran StrandPhaseR^18^ to phase SNVs into whole-chromosome haplotypes. Such sparse haplotypes are then used as a haplotype backbone for long-read phasing using WhatsHap in order to increase density of phased SNVs.

breakpointR command (run and export of results):

~~~
breakpointr(inputfolder = <>, outputfolder = <>, windowsize = 500000,
binMethod = ‘size’, pairedEndReads = TRUE, pair2frgm = FALSE, min.mapq = 10,
filtAlt = TRUE, background = 0.1, minReads = 50)
exportRegions(datapath = <>, file = <>, collapseInversions = TRUE,
collapseRegionSize = 5000000, minRegionSize = 5000000, state = ‘wc’)
~~~

StrandPhaseR command:

~~~
strandPhaseR(inputfolder = <>, positions = <SNVs.vcf>, WCregions =
<hap.informtive.regions>, pairedEndReads = TRUE, min.mapq = 10, min.baseq =
20, num.iterations = 2, translateBases = TRUE, splitPhasedReads = TRUE)
~~~

WhatsHap command:

~~~
whatshap phase --chromosome {chromosome} --reference {reference.fasta}
{input.vcf} {input.bam} {input.vcf_sparse_haplotypes}
~~~

### Haplotagging PacBio reads

Having completely phased chromosomal scaffolds at sufficient SNV density allows us to split long PacBio reads into their respective haplotypes using WhatsHap. This step can be performed in two ways: Splitting all reads across all clusters into two bins per haplotype or splitting reads into two bins per cluster and per haplotype. Both strategies consist of the same two steps: (i) label all reads with their respective haplotype (“haplotagging”) and (ii) splitting the input reads only by haplotype, or by haplotype and cluster (“haplosplitting”). The WhatsHap commands are identical in both cases except for limiting WhatsHap to a specific cluster during haplotagging and discarding reads from other clusters to separate the reads by haplotype and cluster:

~~~
whatshap haplotag [--regions {cluster}] --output {output.bam} --reference
{input.fasta} --output-haplotag-list {output.tags}{input.vcf} {input.bam}
~~~

~~~
whatshap split [--discard-unknown-reads] --pigz --output-h1 {output.hap1} --
output-h2 {output.hap2} --output-untagged {output.un} --read-lengths-
histogram {output.hist} {input.fastq} {input.tags}
~~~

### Creating haplotype-specific assemblies

After haplotagging and -splitting, the long HiFi reads separated by haplotype were then used to create fully haplotype-resolved assemblies. Our haplotagging and -splitting strategy enabled us to examine two types of haploid assemblies per input long-read dataset: the two haplotype-only assemblies (short: h1 and h2), plus the haploid assemblies created by using also all untagged reads, i.e., all reads that could not be assigned to a haplotype (short: h1-un and h2-un). Hence, for each input read dataset, this amounts to four “genome-scale” assemblies. We focused our analyses on the read sets h1-un (H1) and h2-un (H2). Final phased assemblies were created using parameters stated in ‘Collapsed genome assembly’ section.

### SD analysis

SDs were defined as resolved or unresolved based on their alignments to GRCh38 using the minimap2 parameters following parameters: `--secondary=no -a --eqx -Y -x asm20 -m 10000 -z 10000,50 -r 50000 --end-bonus=100 -0 5,56 -E 4,1 □B 5`. Alignments that extended a minimum number of base pairs beyond the annotated SDs were considered to be resolved. The percent of resolved SDs was determined for minimum extension varying from −10,000 to 50,000 bp and average was reported. This analysis is adapted from Vollger et al., 2019 *Nat Meth* (https://github.com/mrvollger/segdupplots).

### Collapse analysis

Collapses were identified as regions in the assemblies that were at least 15 kbp in length and had read coverage exceeding the mean coverage plus three standard deviations. Additionally, collapses that were more than 75% common repeat element (identified with RepeatMasker) or tandem repeats (identified with Tandem Repeats Finder^42^) were excluded.

### BAC clone insert sequencing

BAC clones from the VMRC62 clone library were selected from random regions of the genome not intersecting with an SD (n = 77). DNA from positive clones was isolated, screened for genome location, and prepared for long-insert PacBio sequencing as previously described (SDA)^43^. Libraries were sequenced on the PacBio RS II with P6-C4 chemistry (17 clones) or the PacBio Sequel II with Sequel II 2.0 chemistry (S/P4.1-C2/5.0-8M, 60 clones). We performed *de novo* assembly of pooled BAC inserts using Canu v1.5 (Koren et al., 2017) for the 17 PacBio RS II BACs and using the PacBio SMRT Link v8.0 Microbial assembly pipeline (Falcon + Raptor, https://www.pacb.com/support/software-downloads/) for the 60 Sequel II BACs. After assembly, we removed vector sequence (pCCBAC1), restitched the insert, and then polished with Quiver or Arrow. Canu is specifically designed for assembly with long error-prone reads, whereas Quiver/Arrow is a multi-read consensus algorithm that uses the raw pulse and base call information generated during SMRT sequencing for error correction. We reviewed PacBio assemblies for misassembly by visualizing the read depth of PacBio reads in Parasight (http://eichlerlab.gs.washington.edu/jeff/parasight/index.html), using coverage summaries generated during the resequencing protocol.

### Assembly polishing and error correction

Assembly misjoints are visible using Strand-seq as a recurrent changes in strand state inheritance along a single contig. Strand-state changes result from a double-strand break (DSB) repair during DNA replication and thus observing a strand-state change at the same position in multiple single cells is highly unlikely and therefore indicative of a different process than DSB. Observing a complete switch from WW to CC or vice versa at about 50% frequency is observed when a part of the contig is being misoriented (**Supplementary Fig. 1**). All detected misassemblies in the final phased assemblies were corrected using SaaRclust.

### Common assembly breaks

To detect recurrent breaks in our assemblies we searched for assembly gaps present in at least one phased assembly completed by Canu or Peregrine. For this we mapped all haplotype-specific contigs to GRCh38 using minimap2 using the same parameters as in the SD analysis method. We defined an assembly break as a gap between two subsequent contigs. We searched for reoccurring assembly breaks in 500 kbp non-overlapping bins and filtered out contigs smaller than 100 kbp. Each assembly break was defined as a range between the first and the last breakpoint found in any given genomic bin. Each assembly break was annotated based on the overlap with known SDs, gaps, centromeres and SV callset^19^ allowing overlaps within 10 kbp distance from the breakpoint boundaries.

### Base accuracy

Phred-like QV calculations were made by aligning the final assemblies to 77 sequenced and assembled BACs from VMRC62 falling within unique regions of the genome (>10 kbp away from the closest SD) where at least 95% of the BAC sequence was aligned. The following formula was used to calculate the QV, and insertions and deletions of size N were counted as N errors: QV = –10log10[1 – (percent identity/100)].

Each assembly was polished twice with Racon (Vaser et al., 2017) using the haplotype-partitioned HiFi fastqs. The alignment and polishing steps were run with the following commands:

~~~
minimap2 □ax map□pb --eqx □m 5000 □t {threads} --secondary=no {ref} {fastq}
| samtools view □F 1796 □ > {sam}
racon {fastq} {sam} {ref} □u □t {threads} > {output.fasta}
~~~

QV estimates based on variant call sets lifted back to the human reference hg38 were derived as follows: Genome in a Bottle^44^ high-confidence region sets (release v3.3.2) for individuals HG001, HG002, and HG005 were downloaded and the intersection of all regions (BEDTools v2.29.0 “multiinter”^45^) was used as proxy for high-confidence regions in HG00733 (covering ~2.17 Gbp). HG00733 call sets were generated usingFreeBayes v1.3.1-dirty based on BWA v0.7.17-r1188 alignments of Illumina paired-end short reads (2×125 bp, ~79x coverage) against the haploid HG00733 assemblies:

~~~
bwa mem -t {threads} -R {read_group} {index_prefix} {reads_mate1}
{reads_mate2} | samtools view -b -F 780 - > {output_bam}
~~~

The BAM files were sorted with Samtools v1.9 and duplicates marked with Sambamba v0.6.6 “markdup”. The variant calls with FreeBayes were generated as follows:

~~~
freebayes --use-best-n-alleles 4 --skip-coverage 125 --region
{assembly_contig} -f {assembly_fasta} {input_bam}
~~~

Options “--use-best-n-alleles” and “--skip-coverage” were set following developer recommendations to increase processing speed. Variants were further filtered with BCFtools v1.9: “QUAL >= 10 --genotype hom --max-alleles 2 --types snps,indels”. Variants were converted into BED format using vcf2bed v2.4.37^46^ with parameters “--snvs”, “--insertions”, and “--deletions”. The alignment information for lifting variants from the HG00733 haploid assemblies to the human hg38 reference was generated with minimap v2.17-r941, and the liftover realized with paftools (part of the minimap package):

~~~
minimap2 -t {threads} -c -x asm20 --cs hg38.fasta {input_hap_assembly} >
{hap-assm}_to_hg38.paf
paftools.js liftover -l {min_aln_size} {input_paf} {input_bed} >
{output.hg38.bed}
~~~

To evaluate the robustness of the liftover process, we tested three different values for the minimum alignment size used by paftools: 50 kbp, 25 kbp, and 1 kbp. The lifted variants were intersected with our set of HG00733 high-confidence regions using BEDTools “intersect”. The total number of base pairs in homozygous variants was then computed as the sum over the length (as reported by FreeBayes as LEN) of all variants located in the high-confidence regions. The same process was applied to generate call sets using PacBio HiFi reads of the parent individuals HG00731 (~32x coverage) and HG00732 (~21x coverage), with the following adaptations: initial alignments were generated using minimap v2.17-r941:

~~~
minimap2 -a -x asm20 -R {read_group} -t {threads}
{input_hap_assembly} {input_parent_reads}
~~~

The “--skip-coverage” option of FreeBayes was set to 40 for HG00732, and to 50 for HG00731. The HiFi-based variants were not restricted to the high-confidence regions because long PacBio reads exhibit higher mappability rates throughout the genome compared to short Illumina reads.

### SV and indel detection

Methods for SV and indel calling are similar to previous HiFi assembly work^47^ but were adapted for phased assemblies. Variants were called against the GRCh38 primary assembly (i.e., no alternate, patch, or decoy sequences), which includes chromosomes and unplaced/unlocalized contigs. Mapping was performed with minimap2 2.17^48^ using parameters `“-- secondary=no -a -t 20 --eqx -Y -x asm20 -m 10000 -z 10000,50 -r 50000 --end-bonus=100 -0 5,56 -E 4,1 -B 5”` as described previously^47^. Alignments were then sorted with samtools 1.9^49^.

To obtain variant calls, alignments were processed with PrintGaps.py, which was derived in the SMRT-SV v2 pipeline (https://github.com/EichlerLab/smrtsv2)^50,51^. to parse CIGAR string operations to make variant calls^30^.

Alignment records from assemblies often overlap, which would produce duplicate indel and SV calls with possible different representations (fragmented or shifted). For each haplotype, we constructed a tiling path covering GRCh38 once and traversing loci most centrally located within alignment records. Among the discovered SVs, variants within the path were chosen, and variants outside the tiling path (i.e., potential duplicates) were dropped from further analysis.

After obtaining a callset for haplotype 1 (h1) and haplotype 2 (h2) independently, we then merged the two haplotypes into a single callset. For homozygous calls, an h2 variant must intersect an h1 variant by a) 50% reciprocal overlap (RO), or b) within 200 bp and a 50% reciprocal size overlap (RO if variants were shifted to maximally intersect). The result is a unified phased callset containing homozygous and heterozygous variants. Finally, we filtered out variants in pericentromeric loci where callsets are difficult to reproduce ^51^ and loci where we found a collapse in the assembly of either haplotype.

We intersected RefSeq annotations from the UCSC RefSeq track and evaluated its effect on genes noting frameshift indels in coding regions using custom code to quantify the number of bases affected per variant on genic regions, i.e., exon, intron, and untranslated regions (UTRs).

Variants falling within tandem repeats (TRs) and SDs were also annotated using UCSC hg38 tracks. For TR and SD BED files, we merged records allowing regions within 200 bp to overlap with BEDTools^45^. SVs and indels that were at least 50% contained within an SD or TR region were annotated as SD or TR. For RefSeq analysis, we classified genes as contained within TR or SD by intersecting exons with the collapsed TR and SD regions allowing any overlap.

### MHC analysis

We extracted the MHC, defined as chr6:28000000-34000000, by mapping each haplotype sequence against GRCh38 and extracting any primary or supplementary alignments to this region. We created a dotplot for each haplotype’s MHC region using Dot from DNAnexus (https://github.com/dnanexus/dot) (**Supplementary Figure 11**). We created phased VCFs for both the CCS and Shasta assemblies using the two haplotype files as input to Dipcall (https://github.com/lh3/dipcall). Then, we compared the phasing between the haplotype files using the compare module within WhatsHap. This results in a switch error rate of 0.48% (6 sites) and Hamming error rate of 0.28% (4 sites) from 1,433 common heterozygous sites between the vcfs.

#### Likely disrupted genes

Using RefSeq intersect counts, we found all genes with at least one non-modulo-3 insertion or deletion within the coding region of any isoform (i.e., frameshift). We filtered out any genes not fully contained within a consensus region of the two haplotypes, which we defined as regions where both h1 and h2 had exactly one aligned contig. If a gene had multiple non-modulo-3 events, whether in the same isoform or not, the gene was counted once.

### Variant comparisons

We compared variants to previously published callsets by intersecting them with the same RO/Size-RO strategy that was used to merge haplotypes. For HGSVC comparisons, we also excluded variant calls on unplaced contigs, unlocalized contigs, and chrY of the reference (i.e., chr1-22,X), which were not reported by the HGSVC study. To quantify the number of missed variants proximal to another, we took variants that failed to intersect an HGSVC variant and found the distance to the nearest variant of the same type (INS vs. INS and DEL vs. DEL).

### Robust and reproducible implementation

The basic workflow of our study is implemented in a reproducible and scalable Snakemake^52^ pipeline that has been successfully tested in compute environments ranging from single servers to high-performance cluster setups (**Code availability**). Major tasks in the pipeline, such as read alignment or assembly, have been designed as self-contained “start-to-finish” jobs, automating even trivial steps such as downloading the publicly available datasets used in this study. Due to the considerable size of the input data, we strongly recommend deploying this pipeline only on compute infrastructure tailored to resource-intensive and highly parallel workloads.

## Code availability

R package SaaRclust (MIT License): https://github.com/daewoooo/SaaRclust (devel branch)

R package breakpointR (MIT License): https://bioconductor.org/packages/breakpointR/

R package StrandPhaseR (MIT License): https://github.com/daewoooo/StrandPhaseR (devel branch)

Snakemake pipeline (MIT License): https://github.com/ptrebert/project-diploid-assembly (development branch)

## Data availability

HiFi PacBio reads for HG00731, HG00732, and HG00733 were produced as part of this study and are available from the IGSR FTP (ftp.1000genomes.ebi.ac.uk/vol1/ftp/data_collections/HGSVC2/working/20190925_PUR_PacBio_HiFi/) and will be accessioned on EBI for print. Strand-seq data for HG00733 were downloaded from NCBI SRA (BioProject PRJEB12849). Illumina short reads for HG00733 were downloaded from NCBI SRA (BioProject PRJEB9396). The genome assemblies produced in this study are available from http://ftp.1000genomes.ebi.ac.uk/vol1/ftp/data_collections/HGSVC2/working/20191122_Marschall-Eichler_HG00733_HiFi_hap-assm. We make available two versions of the assembly: “Freeze 1” and “Freeze 1.1”. The later Freeze 1.1 is based on exactly the same pipeline, but a minor software bug (in the SaaRclust module) was fixed that led to some contig misorients of the collapsed assembly resulting in phasing inconsistencies downstream. All statistics are based on Freeze 1, unless indicated otherwise.

## Acknowledgements

The authors thank T. Brown for assistance in editing this manuscript, Chen-Shan Chin for advice on setting optimal parameters for Peregrine assembly and the Human Genome Sequencing Consortium (HGSVC) for data access and comments. A full list of HGSVC contributors is found in the supplementary material.

## Author contributions

D.P., P.E., E.E.E, and T.M. designed the study. P.E. and D.P. implemented the assembly workflow. P.A.A. and M.J.P.C. performed structural variant analysis. M.R.V. and D.P. analyzed assemblies for universal breaks, segmental duplications and collapses. An earlier HiFi data set was provided by S.E.D. and used during method development. M.H. and B.P. compared assemblies to trio-binned Shasta assemblies. HGSVC members engaged in fruitful discussions, led by C.L., at the biannual consortium meetings. W.T.H performed variant calling for phasing. K.M.M generated HiFi PacBio data. M.S. sequenced BAC clones for validation. A.S. analyzed tandem repeat variants. D.P., P.A.A., M.R.V., T.M. prepared main display items. D.P., P.E., P.A.A., M.R.V., E.E.E., and T.M. wrote the manuscript, with input from A.D.S., M.G., P.M.L, and J.O.K.

