## Supplementary Information for "A fully phased accurate assembly of an individual human genome"

### Supplementary notes

#### Strand-seq

The Strand-seq protocol utilizes a thymidine analog to selectively label and remove one of the DNA strands (the nascent strand, synthesized during DNA replication) and leaves only the template DNA strands intact for sequencing. By sequencing only template strands in each homologue, Strand-seq distinguishes three possible template strand states for each chromosome of a diploid genome. The Watson-Watson (WW) strand state is characteristic of two Watson (reads aligned to minus strand) templates inherited from both parental homologues. The Crick-Crick (CC) strand state is characteristic of two Crick (reads aligned to plus strand) templates inherited from both parental homologues. Lastly, the Watson-Crick (WC) strand state is characteristic of a Watson and Crick template being inherited from either parental homologue. In this WC scenario the two parental templates can be distinguished based on read directionality and are thus informative of phasing. Template strands are randomly inherited by single daughter cells, resulting in a specific strand-state pattern for each chromosome across multiple Strand-seq libraries (**Fig. 1b**). This strand-state pattern can be viewed as a barcode that uniquely assigns each contig to its chromosome of origin (Hills et al. 2013) (**Fig. 1c**). However, we do not always observe a single strand state along the whole chromosome but instead there can be strand-state changes as a result of double-strand break (DSB) repaired by sister chromatid exchange (SCE) during DNA replication (Falconer et al. 2012; van Wietmarschen and Lansdorp 2016; Claussin et al. 2017). Such low frequency SCEs are indicative of the physical distance between two segments of a chromosome, because segments that are physically further apart from each other have an increased likelihood of an SCE occurring between them (Hills et al. 2013). This means that contigs that are physically linked to each other are less likely to be separated by SCEs and thus will share the same strand state across multiple cells—a signal that enables assembled contigs to be clustered into chromosomes and then ordered within each chromosome (**Fig. 1d**). Clustered and ordered contigs can then be phased using single-nucleotide polymorphism information extracted from the haplotype-informative (WC) regions in the Strand-seq data (**Fig. 1e**). This allows us to physically separate parental alleles along the whole chromosome (Porubský et al. 2016). Such global phasing information in conjunction with long-read technologies such as PacBio allows us to reconstruct highly accurate and nearly complete haplotypes that span the whole chromosomes (Porubsky et al. 2017; Chaisson et al. 2019). Such haplotypes serve as a guide to divide long-read data into two bins, one for each haplotype (**Fig. 1f**).

### SaaRclust

Every chromosome undergoes independent random segregation during cell division, leading to a unique strand-state profile in Strand-seq data. This signal in Strand-seq data can be employed to cluster long sequencing reads by chromosome of origin and sequencing direction. SaaRclust (Ghareghani et al. 2018) is a tool that we previously introduced for this *in silico* separation of long reads by chromosome and direction. SaaRclust employs an Expectation-Maximization (EM) soft clustering algorithm to handle the uncertainty arising from the sparse Strand-seq data. Given the central importance of SaaRclust for the assembly pipeline we introduce here, we include Supplementary Figure 12 illustrating the principle. The main idea underlying our clustering algorithm is that contigs originating from the same chromosome (Contigs 1 and 2 in Supplementary Fig. 12) show the same directionality pattern of aligned Strand-seq reads across single cells, which is different for contigs originating from different chromosomes (Contig 3 in Supplementary Fig. 12). The EM algorithm is based on iterating between assigning strand states for each Strand-seq library and chromosome and assigning chromosomes to each contig, which are both hidden information at the beginning. EM converges to a local optimum solution of the maximum likelihood problem, e.g., maximizing the likelihood of observed data (number of directional aligned Strand-seq reads to long reads), given the model parameters (strand states), and we have shown SaaRclust to be able to assign even individual long reads to chromosomes of origin. Here, we have adjusted it to work on the contig level.

### Variant discovery and comparisons

The SNV transition/transversion (Ti/Tv) proportion was 1.99, 1.98, and 1.98 for h1, h2, and merged callset, respectively. Outside of tandem repeats, the Ti/Tv rose to 2.05.

For variants that did not intersect an HGSVC call, we find that 78% (3,079 of 3,945) of false insertions and 75% (1,527 of 2,048) of false deletions map within 1 kbp of a variant of the same type indicating that many of these calls may be different representations of the same event but represent inconsistent alignment as discussed previously. Squashed assemblies, even when reference-guided, miss a large proportion heterozygous SV calls (Huddleston et al. 2017), and compared to a haplotype-unaware analysis of HG00733 (Audano et al. 2019), we find 31% and 12% more insertions and deletions outside repetitive loci, respectively.

### Supplementary Figures 1-12

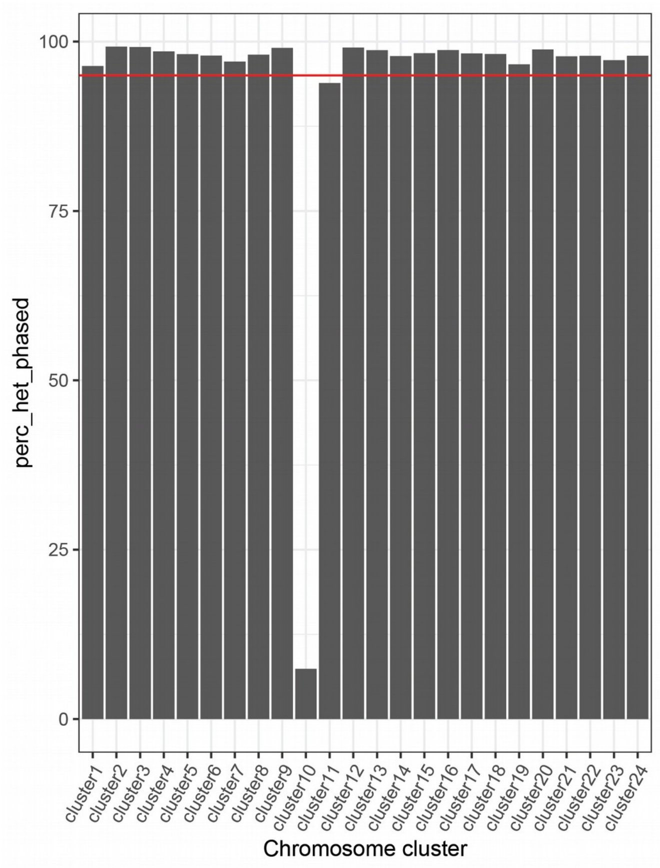


**Supplementary Figure 1: Phasing of SNVs per chromosomal cluster.**
Height of each bar represents the percentage of SNVs phased in the longest haplotype block in each cluster. Red line highlights the 95% threshold.

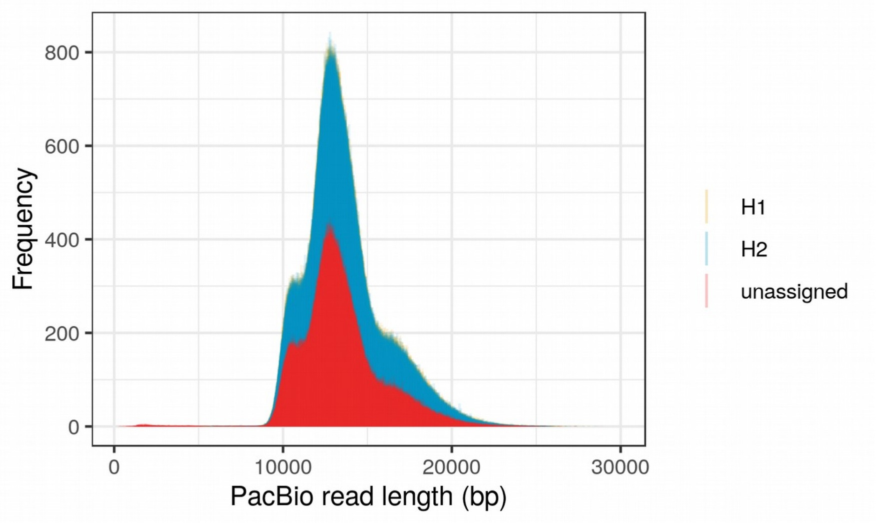


**Supplementary Figure 2: Size distribution of haplotagged reads.**
Distributions indicate observation frequencies (y-axis) of different read lengths (x-axis) for the HG00733 HiFi dataset split by haplotype (“haplotagged”). The two haplotype fractions H1 and H2 are plotted as yellow and green distributions, respectively. The fraction of unassigned (“untagged”) reads is plotted in red.

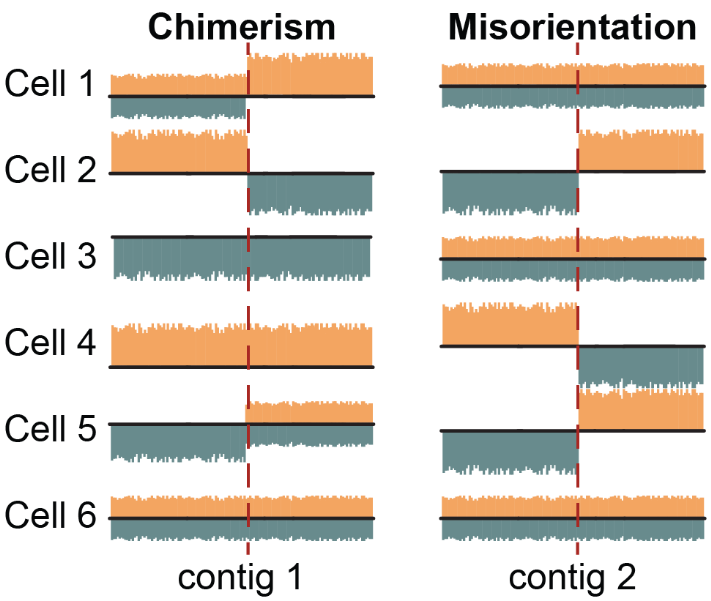


**Supplementary Figure 3: Strand-seq patterns of common misassemblies.**
A genome misassembly is visible in Strand-seq data as a recurrent change in strand state at the same position in a given contig. Because, it is highly unlikely for a double-strand break to occur at exactly the same position in multiple single cells, the most likely explanation in this case is either contig misorientation or chimerism. Chimerism is characteristic by almost all possible template strand changes as the portions of a chimeric contig carry a strand state of the contig they truly belong to. On the other hand, misorientation is characteristic of a complete switch from either WW to CC or vice versa. This type of misassembly is visible in about 50% of cells as only WW or CC template strand states are informative for this type of assembly error.

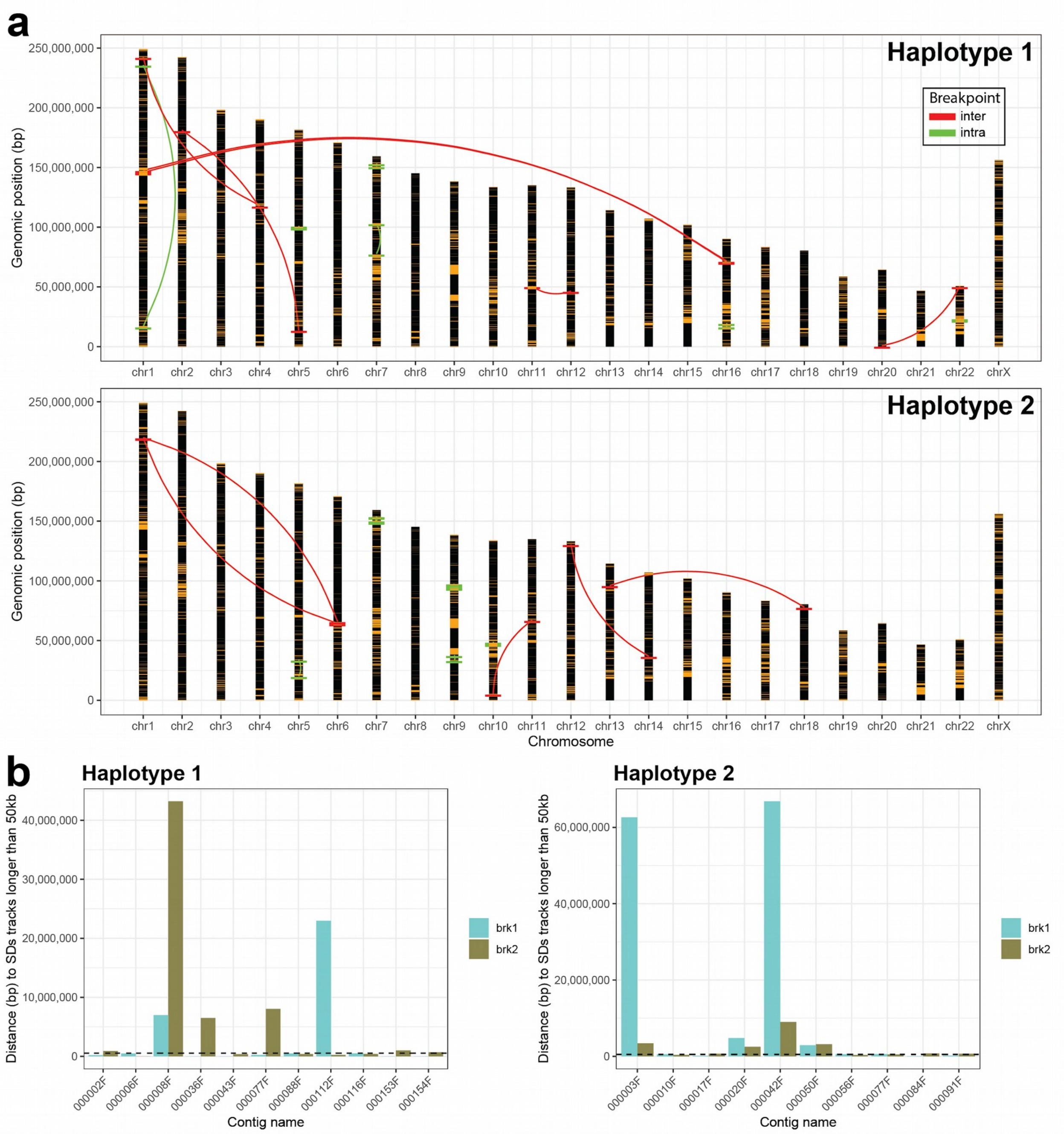


#### Supplementary Figure 4: Peregrine-specific assembly errors.

**a**) Projection of Peregrine-based assembly errors to GRCh38. Positions of segmental duplications (SDs) in the genome are highlighted in orange. Red and green links connect regions upstream and downstream from an assembly error (**Methods**). If no link is visible, position upstream and downstream from the breakpoint lies in close proximity. **b**) Each bar (turquoise - upstream from the assembly error, khaki - downstream from the assembly error) represent a distance to the closest SD track of 50 kbp and longer from the assembly error (turquoise - upstream from the assembly error, khaki - downstream from the assembly error) per misassembled contig (x-axis).

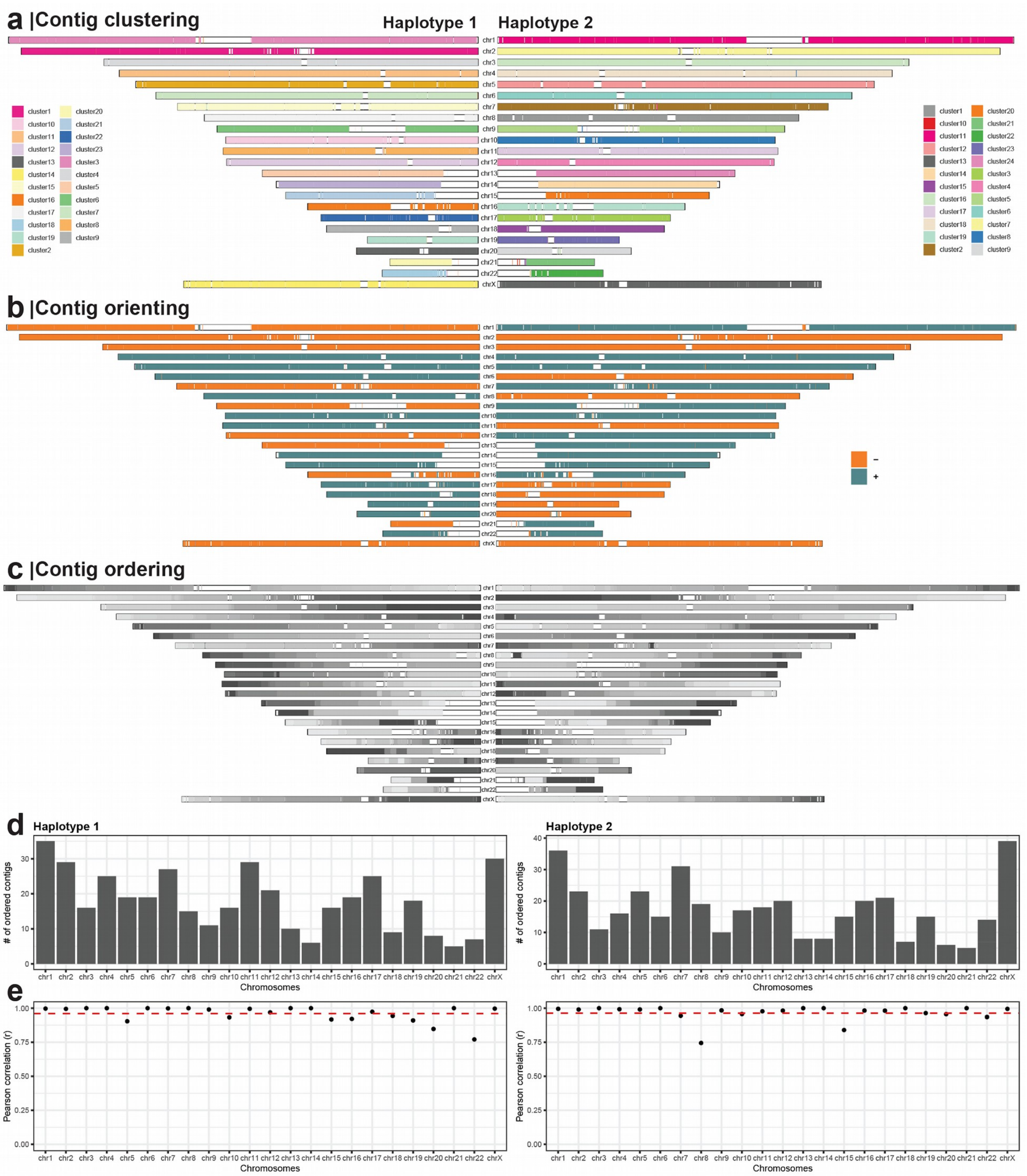


**Supplementary Figure 5: Accuracy of contig ordering within each chromosomal cluster.** For this analysis we used only contigs 500 kbp and longer and those that can be assigned to a chromosomal cluster with probability p >= 0.9. **a**) Each contig represents a range based on mapping coordinates on GRCh38. Contigs are colored based on cluster identity determined by SaaRclust. In an ideal scenario there is a single color for each chromosome. **b**) Each contig represents a range based on mapping coordinates on GRCh38. Contigs are colored based on the directionality ('+' - positive strand, '-' - negative strand) they map to GRCh38. In an ideal scenario there is a single color for each chromosome. **c**) Each contig is colored based on the predicted order within each chromosomal cluster which is reflected by the shades of gray going from dark to light gray. Ideally we observe colors going always from dark to light gray or vice versa and thus being in agreement with true contig order on GRCh38. **d**) Each bar represents a number of contigs submitted for ordering within each chromosome and haplotype. **e**)

Correlation of predicted contig order with the expected ordering, based on GRCh38 mappings, within each chromosome and haplotype. Red dashed line shows mean correlation over all chromosome within a haplotype.

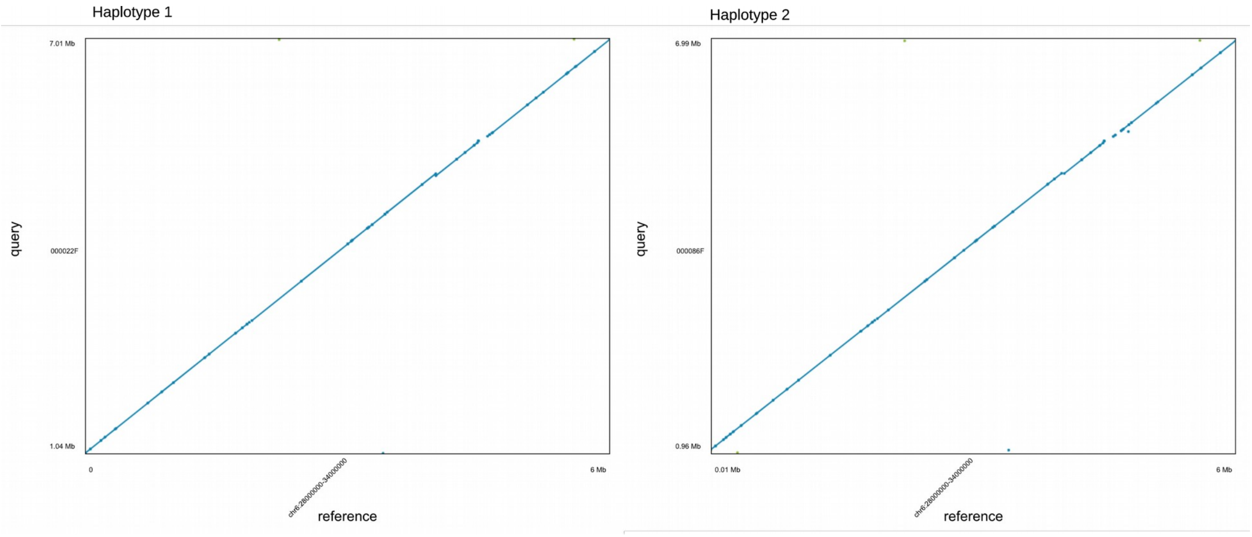


#### Supplementary Figure 6: Dot plots of the major histocompatibility complex (MHC).

In each haplotype assembly, the whole MHC region is traversed by one single contig. Above we show a corresponding dot plot of the two haplotypes H1 (left) and H2 (right).

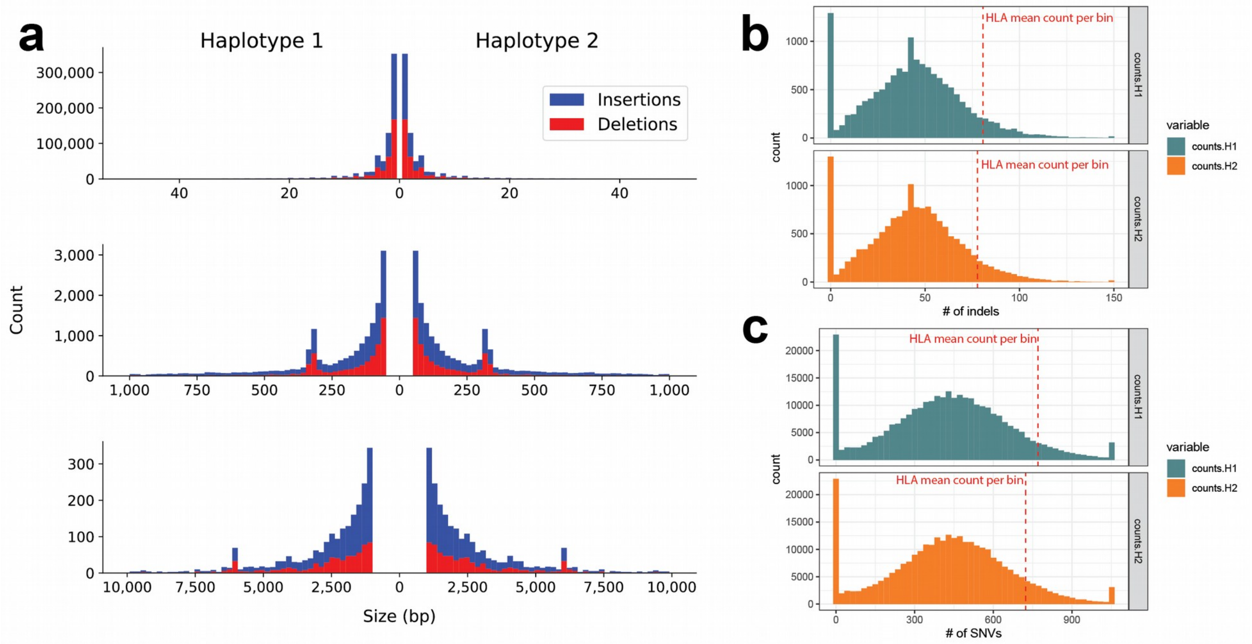


#### Supplementary Figure 7: Indels density in phased assemblies.

**a**) Top: Single base-pair indels (1-49 bp) are most common with peaks modulo-2 bp events (4, 6, 8, etc.) corresponding to the prevalence of dinucleotide repeat elements. Middle: Smaller SVs (50-999 bp) show a peak for SINE elements. Bottom: Larger SVs show a peak for LINE elements. SVs larger than 10,000 are not shown. **b**) and **c**) Histograms showing the distribution of small indels in (b) and SNVs in (c) counted in 200 kbp long non-overlapping bins separately for haplotype 1 (H1 - teal) and haplotype 2 (H2 - orange). Mean indel and SNV count in bins spanning the HLA locus is highlighted by a red dashed lines. Indels and SNVs in regions of detected assembly collapses and known SDs have been removed.

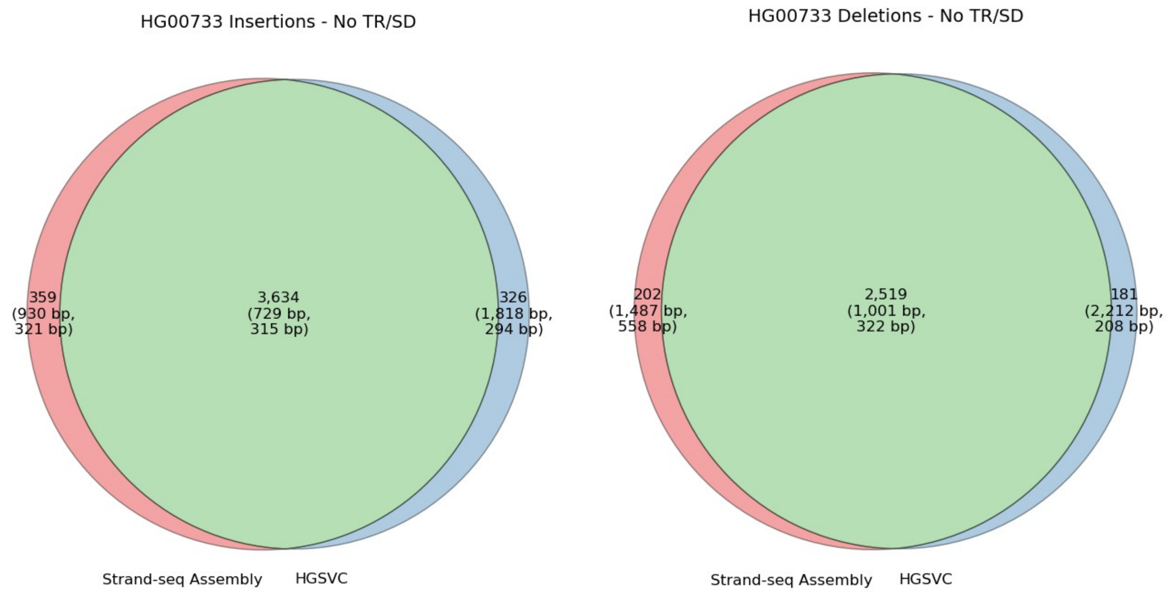


**Supplementary Figure 8: Variant comparisons between assembly SVs and HGSVC HG00733 outside tandem repeat (TR) and segmental duplication (SD) loci.**
Variant comparisons outside TRs and SDs give a picture of concordance without many of the alignment problems that make repeats difficult to represent and reproduce. The number of variants is shown with the mean (top) and median (bottom) SV size in parentheses. Unplaced and unlocalized SVs were removed from this analysis, which were filtered in HGSVC.

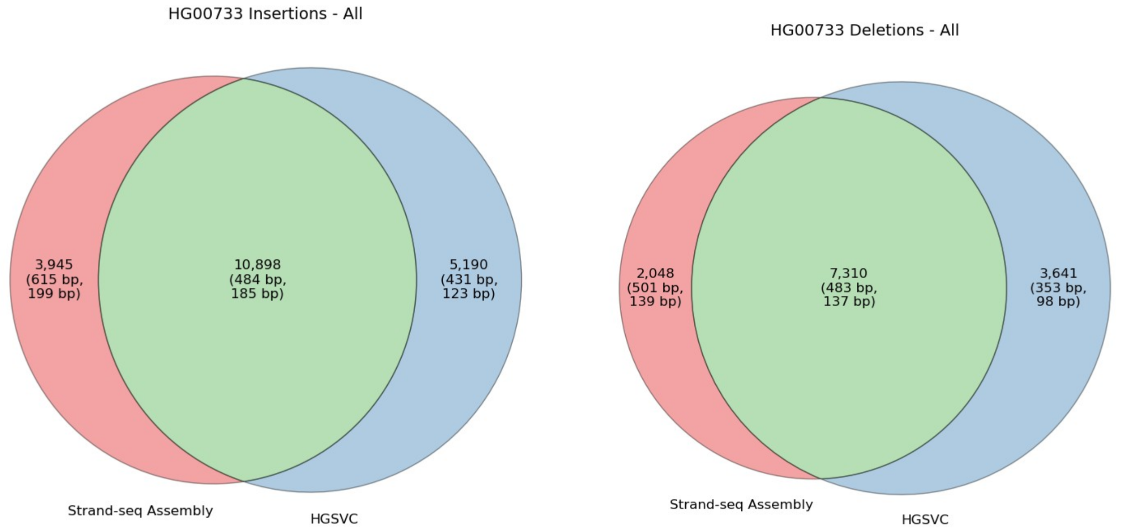


**Supplementary Figure 9: Variant comparisons between assembly SVs and HGSVC HG00733.**
Variant comparisons including TRs and SDs are harder to replicate, even for larger events, which are often fragmented or shifted by alignments through repeats. The number of variants is shown with the mean (top) and median (bottom) SV size in parentheses. Unplaced and unlocalized SVs were removed from this analysis, which were filtered in HGSVC.

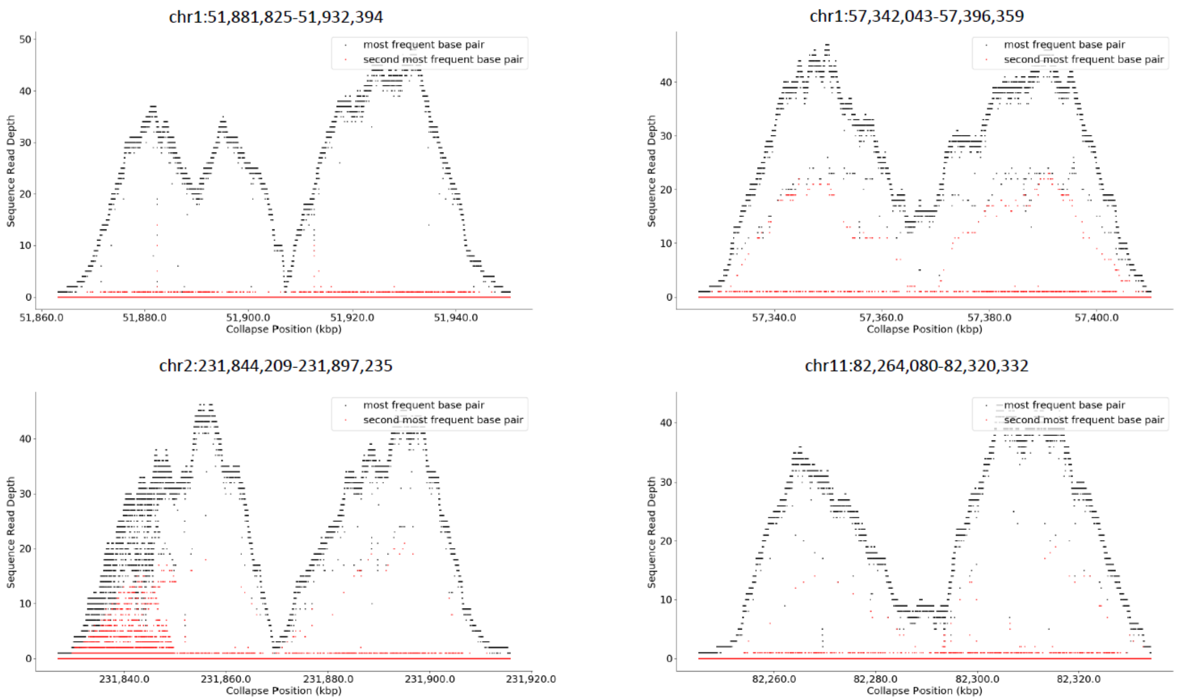


#### Supplementary Figure 10: SV signatures at assembly breakpoints.

Shown are coverage profiles of HiFi reads aligned to GRCh38 at four different assembly breakpoints. The coverage for the most frequent and for the second most frequent base at each position is shown in black and red, respectively. These coverage profiles are consistent with structural variation at these locations.

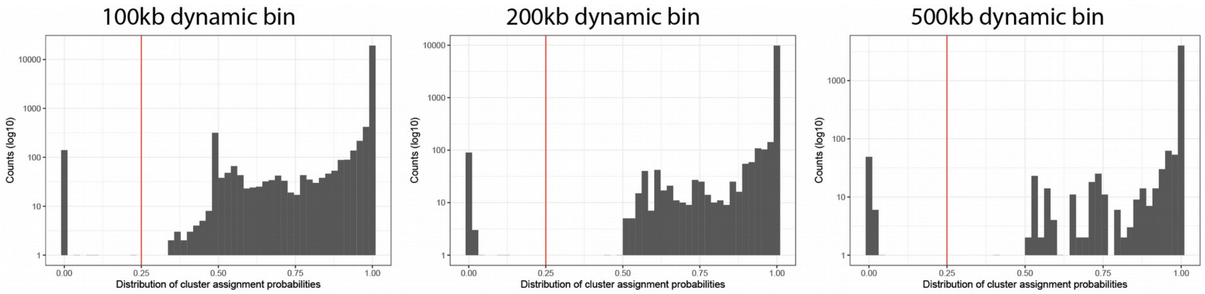


#### Supplementary Figure 11: Effect of binning strategy on final cluster assignment probabilities.

In SaaRclust each piece of DNA (contig) is assigned a probability of belonging to any of the tested chromosomal clusters. This probability is calculated using the previously published expectation maximization algorithm (Ghareghani et al. 2018). As expected, some contigs are difficult to assign unambiguously to a single cluster and could be assigned to several clusters with equally low probabilities (left part of the distribution in panels). We examined the effect of varying the bin size from 100 kbp to 500 kbp on the resulting probability distribution. Given the observed probability distribution, we decided to set a dynamic bin size to 200 kbp (SaaRclust ‘bin.size’ parameter) with the probability threshold (SaaRclust ‘prob.th’ parameter) set to 0.25.

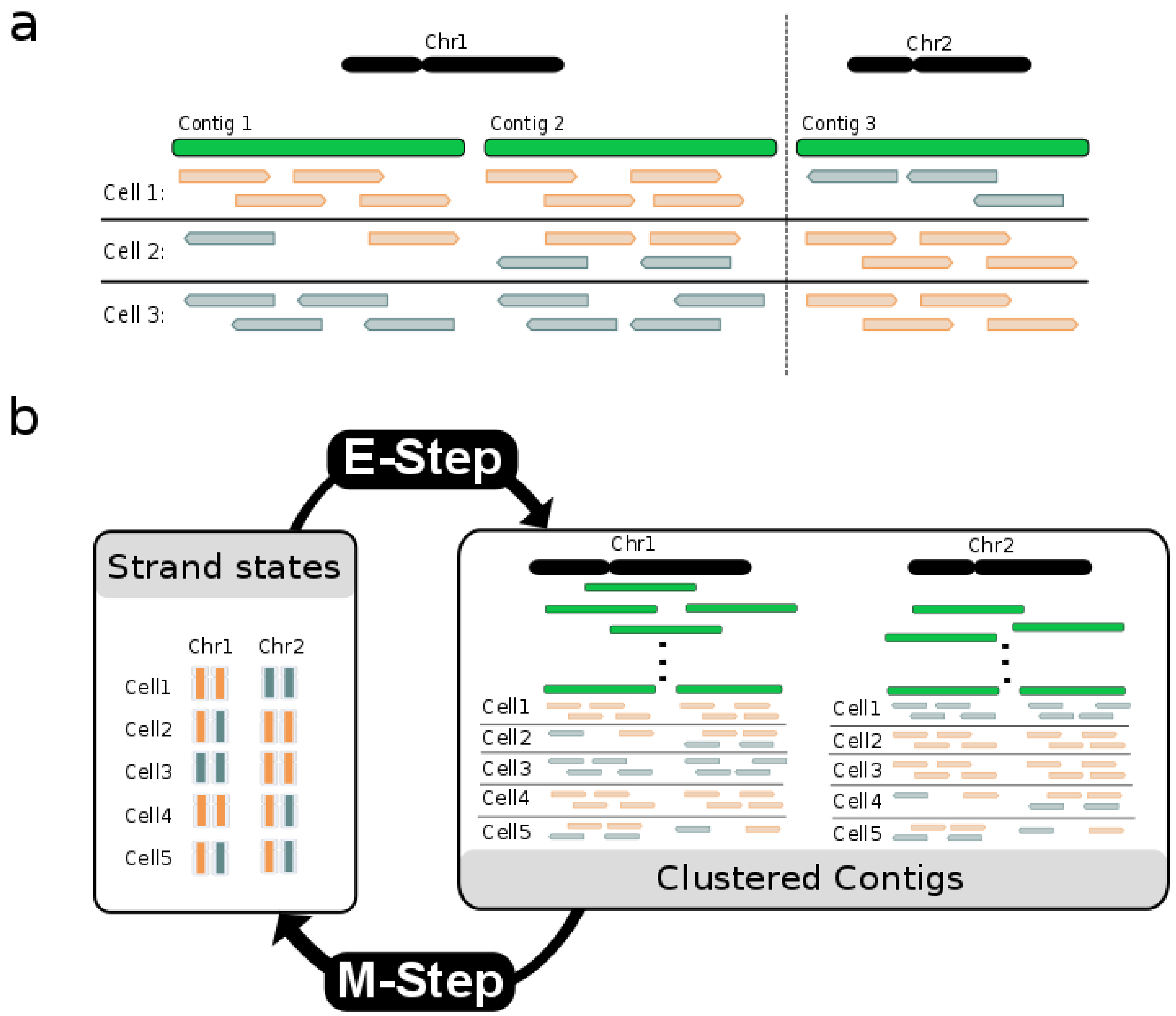


**Supplementary Figure 12: An overview of SaaRclust approach for clustering contigs by chromosome and orientation** (Ghareghani et al. 2018)**.**

**a**) Aligning single-cell Strand-seq reads to collapsed contigs. In this example, Strand-seq reads from three different single cells are aligned to three contigs. Strand-seq reads mapped in Watson and Crick directions to contigs are shown by orange and teal colors, respectively. Contigs 1 and 2 come from Chromosome 1 showing a different strand state from Contig 3 that comes from Chromosome 2. **b**) Schematic of the EM clustering algorithm for two chromosomes. Starting from an arbitrary initialization of strand states, the EM algorithm iterates through the flow of information between the two hidden layers of information: the strand states of single cells in chromosomes (left box) and the clustering of contigs into chromosomes (right box).

### Supplementary Tables 1-7

#### Supplementary Table 1: *De novo* assembly statistics.

| | Total.size | Total.contigs | N50 (bp) | Asm.errors |
|---|---|---|---|---|
| **canu phased H1** | 3.09Gb | 4844 | 9873595 | 3 chimerisms, 6 misorients |
| **canu phased H2** | 3.08Gb | 4830 | 10725871 | 1 chimerisms, 4 misorients |
| **peregrine phased H1** | 2.9Gb | 2557 | 28001683 | 6 chimerisms, 5 misorients |
| **peregrine phased H2** | 2.91Gb | 2618 | 29160574 | 5 chimerisms, 5 misorients |

#### Supplementary Tables 2A/2B: Variant call-based QV estimates (external xlsx files).

**2A** Illumina short reads (HG00733) were used to call variants (SNVs and indels) relative to the haploid assemblies of HG00733 (top); number of homozygous ("hom") calls listed in detail, total number of heterozygous ("het") calls stated for comparison. Homozygous calls were lifted to human reference hg38 using three minimum alignment sizes (middle left) and restricted to high-confidence regions ("hc", middle right). Percentages relative to all homozygous variants called, and to all homozygous variants lifted to hg38, respectively. Assembly accuracy estimates (QV) were calculated by considering all base pairs ("bp") in homozygous variants as errors (bottom). **2B** PacBio HiFi reads (HG00731 and HG00732) were used to call variants (SNVs and indels) relative to the haploid assemblies of HG00733 (same as in 2A). Since long PacBio reads show higher mappability throughout the genome compared to short Illumina reads, hg38-lifted variants were not restricted to high-confidence regions (middle). QV estimates were calculated as in 2A.

#### Supplementary Table 3: Phased assembly indel discovery.

Indels were discovered in both haplotypes and merged into a single call. Fields are number of variants ("N"), mean indel size ("Mean (bp)"), total number of indel bases ("Base (kbp)"), the percentage 1 bp indels ("1 bp (%)"), and the percentage heterozygous calls ("Het (%)").

| | Insertions | | | | | Deletions | | | | |
|---|---|---|---|---|---|---|---|---|---|---|
| Assembly | N | Mean (bp) | Base (kbp) | 1 bp (%) | Het (%) | N | Mean (bp) | Base (kbp) | 1 bp (%) | Het (%) |
| HG00733 (Racon x2) | 515,224 | 3.44 | 1,773 | 49.97% | 61.29% | 494,810 | 3.71 | 1,835 | 47.23% | 61.93% |
| HG00733 (unpolished) | 514,949 | 3.44 | 1,770 | 50.42% | 61.83% | 530,300 | 3.58 | 1,897 | 49.31% | 62.61% |

#### Supplementary Table 4: Phased assembly structural variation discovery.

Variants were discovered in both haplotypes and merged to a set of homozygous and heterozygous calls. Fields are number of variants ("N"), mean variant size ("Mean (bp)"), sum of all variant lengths ("Base (Mbp)"), and the percentage of heterozygous calls by number ("Het (N%)") and by bases ("Het (bp %)").

| | Insertions | | | | | Deletions | | | | |
|---|---|---|---|---|---|---|---|---|---|---|
| Assembly | N | Mean (bp) | Base (Mbp) | Het (N%) | Het (bp %) | N | Mean (bp) | Tput (Mbp) | Het (N%) | Het (bp %) |
| HG00733 (Racon x2) | 15,139 | 521 | 7.89 | 58.71% | 51.99% | 9,579 | 490 | 4.69 | 65.55% | 65.82% |
| HG00733 (unpolished) | 15,174 | 519 | 7.88 | 58.81% | 52.45% | 9,545 | 491 | 4.68 | 66.38% | 66.74% |

#### Supplementary Table 5: Frameshift-disrupted RefSeq annotations.

We quantified the number of genes with a frameshift indel or SV in coding regions and demonstrate that polishing is still required for phased Peregrine assemblies. Shown are disrupted gene counts for all genes ("All"), genes with no exons intersecting tandem repeats or segmental duplications ("No TR/SD"), and genes with at least one exon in a known segmental duplication ("In SD").

| Sample | Assembler | Polishing | All | No TR/SD | In SD |
|---|---|---|---|---|---|
| HG00733 | Peregrine | Racon x2 | 198 | 92 | 42 |
| HG00733 | Peregrine | None | 300 | 111 | 106 |

#### Supplementary Table 6: List of detected universal assembly breaks (external xlsx file).

#### Supplementary Table 7: HiFi PacBio sequencing summary.

| **Sample** | **HG00731** | **HG00732** | **HG00733** |
|---|---|---|---|
| **# SMRT Cell 8Ms** | 5 | 6 | 7 |
| **Raw Base Yield (Gbp)** | 1612 | 1138 | 1568 |
| **HiFi Base Yield (Gbp)** | 103 | 67 | 104 |
| **HiFi Coverage (X)** | 32 | 21 | 32 |
| **Average HiFi Read Length (kbp)** | 11.1 | 10.7 | 13.6 |
| **Median HiFi QV** | 31.86 | 31.59 | 30.39 |
| **Average HiFi number of passes** | 10.51 | 10.54 | 9.34 |

**Human Genome Structural Variation Consortium**

**Steering Committee:** Evan E. Eichler, Jan O. Korbel, Charles Lee
**Consortium Members (alphabetical order):** Haley Abel, Alexej Abyzov, Can Alkan, Thomas Anantharaman, Danny Antaki, Peter A. Audano, Ali Bashir, Mark Batzer, Harrison Brand, Lisa Brooks, Stuart Cantsilieris, Han Cao, Eliza Cerveira, Mark J. P. Chaisson, Ken Chen, Chong Chen, Xintong Chen, Chen-Shan Chin, Zechen Chong, Nelson T. Chuang, Deanna M. Church, Laura Clarke, Ryan L. Collins, Robel Dagnow, Scott E. Devine, Li Ding, Peter Ebert, Susan Fairley, Xian Fan, Andrew Farrell, Ian Fiddes, Paul Flicek, Joey Flores, Daniel Fordham, Timur Galeev, Eugene J. Gardner, Mark B. Gerstein, David U. Gorkin, Madhusudan Gujral, Li Guo, Gamze Gursoy, Victor Guryev, Ira Hall, Robert E. Handsaker, Eoghan Harrington, William Harvey, Alex R. Hastie, William Haynes Heaton, Wolfram Hoeps, Fereydoun Hormozdiari, Junie Jen, Goo Jun, Chong Lek Koh, Xiangmeng Kong, Miriam Konkel, Jonas Korlach, Zev N. Kronenberg, Sushant Kumar, Pui-Yan Kwok, Jee Young Kwon, Sofia Kyriazopoulou-Panagiotopoulou, Ernest T. Lam, Christine C. Lambert, Peter M. Lansdorp, Jong Eun Lee, Sau Peng Lee, Wan-Ping Lee, Dillon Lee, Joyce Lee, Shantao Li, Ernesto Lowy Gallego, Shamoni Maheshwari, Ankit Malhotra, Patrick Marks, Tobias Marschall, Gabor T. Marth, Alvaro Martinez Barrio, Adam Mattson, Steven McCarroll, Sascha Meiers, Ryan E. Mills, Katherine M. Munson, Fabio C. P. Navarro, Bradley J. Nelson, Conor Nodzak, Amina Noor, Andy W. C. Pang, David Porubsky, Letu Qingge, Yunjiang Qiu, Tobias Rausch, Allison Regier, Bing Ren, Oscar L. Rodriguez, Gabriel Rosanio, Joel Rozowsky, Mallory Ryan, Ashley D. Sanders, Michael Schnall-Levin, Jonathan Sebat, Omar Shanta, Steve Sherry, Xinghua Shi, Laura Carolyn Smith, Mike Smith, Diana C. J. Spierings, Adrian Stütz, Arvis Sulovari, Michael E. Talkowski, Karine Viaud-Martinez, Alistair Ward, Anne Marie E. Welch, Jia Wen, Aaron M. Wenger, Matthew Wyczalkowski, Ming Xiao, Wei Xu, Sergei Yakneen, Xiaofei Yang, Kai Ye, Christopher Yoon, Chengsheng Zhang, Xuefang Zhao, Xiangqun Zheng-Bradley, Arthur Zhou, Qihui Zhu, Mike Zody
